# Deciphering gain-of-function from loss-of-function variants with AlphaMissense: A case study with the mechanosensitive PIEZO1 ion channel protein

**DOI:** 10.1101/2025.07.03.662957

**Authors:** Joshua Pillai, Adhvaith Sridhar, Kijung Sung, Linda Shi, Chengbiao Wu

**Affiliations:** School of Biological Sciences, University of California, San Diego, La Jolla, CA, USA; Department of Neurosciences, University of California San Diego, School of Medicine, La Jolla, CA, USA; Biophotonics Technology Center, Institute of Engineering in Medicine, University of California, San Diego, La Jolla, CA, USA; Department of Biochemistry, Molecular Biology, and Biophysics, University of Minnesota Twin Cities, Minneapolis, MN, USA

**Keywords:** Piezo1, Congenital lymphatic dysplasia, Hereditary xerocytosis, AlphaMissense, Mechanosensitive cation channel, Gain-of-Function mutations

## Abstract

Pathogenic missense mutations are commonly found in protein-coding regions of DNA, and often alter protein function. In the past decade, enormous experimental efforts have been undertaken through the development of functional assays, mutagenesis screenings, and computational prediction algorithms to characterize these variants. Indeed, most efforts have been focused towards identifying degrees of loss-of-function (LoF) effects on the three-dimensional structures of proteins, but gain-of-function (GoF) mutations remain poorly understood. Herein, we performed a case study of the PIEZO1 mechanosensitive ion channel protein whose GoF variants (*n* = 56) are implicated in hereditary xerocytosis (HX) and LoF variants (*n* = 6) in lymphatic dysplasia (LD), respectively. This study evaluated the abilities of AlphaMissense (AM) to decipher both mutation types, and benchmarked its performance against other algorithmic approaches, including Combined Annotation Dependent Depletion (CADD) v1.7, evolutionary model of variant effect (EVE), and Evolutionary Scale Modeling-1b (ESM-1B). We found that all approaches excelled in identifying LoF variants but were often ambiguous in their predictions for GoF PIEZO1 variants. ESM-1b was a notable exception that demonstrated balance sensitivity to both GoF and LoF variants as it likely identified certain sequential features not utilized in other approaches. Secondly, our findings suggest that GoF variants of HX do not significantly destabilize the PIEZO1 structure and are not generally identified by conventional signatures of damage from these algorithms. GoF variants are not synonymous with direct changes in changes in free energy upon mutations. Furthermore, we validated computational structure predictions of PIEZO1 against resolved cryo-EM structures, and provided biophysical data for variants located on unresolved residues of this ion channel. We formulated a weighted ensemble model that performed similarly to AM and outperformed all other traditional approaches evaluated in this study. Overall, this is the first study to directly evaluate the capabilities of pathogenicity prediction algorithms for deciphering GoF and LoF variants, and underscores the limitations present in current approaches.

## 1. Introduction

PIEZO1 is a mechanosensitive ion channel comprised of over 2,500 amino acids and 24-36 transmembrane domains forming a homotrimeric propellor structure (Coste et al., 2010; Xiao et al., 2024). PIEZO1 is involved in regulating membrane potential and Ca^2+^ signaling coupled to downstream effectors in animals (Beech et al., 2018). Previous studies have identified that PIEZO1 is an inherent sensor of membranophone tension and a primary physiological activator (Lewis & Grandl., 2015; Cox et al., 2016; Syeda et al., 2016; Beech et al., 2018). PIEZO1 is expressed in numerous cell-types with a wide range in function, including regulating blood pressure, mediating endothelial shear-stress and vascular development, and erythrocyte function (Gudipaty et al., 2017; Zarychanski et al., 2012; Rode et al., 2017; Beech et al., 2018). Further, mutations in PIEZO1 have been implicated in lymphatic dysplasia (LD), hereditary xerocytosis (HX), and other rare disorders (Alper., 2017). Specifically, loss-of-function (LoF) mutations in the *PIEZO1* gene are implicated in autosomal recessive generalized lymphatic dysplasia (Fotiou et al., 2018; Marouli et al., 2017; Lukacs et al., 2015). Compound heterozygous and cosegregating homozygous mutations, including splice-site, nonsense, and missense variants, have been reported in patients (Alper., 2017). In erythrocytes, PIEZO1 mediated Ca^2+^ signaling in response to various mechanical stimuli in heterozygous parents but not in compound heterozygotes of the G2029R variant (Lukacs et al., 2015). Clinical phenotypes observed in PIEZO1-associated lymphatic dysplasia appear distinct from other lymphedema syndromes, and it has been previously suggested that the *PIEZO1* gene may be involved in modulating the expression of other genes implicated in lymphatic endothelial cell signaling, such as *VEGF-C* and *VEGFR-3* (Ma et al., 2008; Alper., 2017). On the other hand, gain-of-function (GoF) mutations in the *PIEZO1* gene have been found in HX from numerous studies (Picard et al., 2019; Russo et al., 2018; Albuisson et al., 2013; Andolfo et al., 2013; Risinger et al., 2017). In GoF mutations, the autosomal dominant missense variants demonstrate altered ion channel kinetics and delayed inactivation (Bae et al., 2013; Alper., 2017). However, the mechanistic relation between the persistent lymphatic insufficiency of LoF and the apparent insufficiency of GoF PIEZO1 variants remains uncertain.

Biochemical and biophysical experimentation with PIEZO1 remains difficult due to the large size and complexity of PIEZO1. Computing restraints on single protein chains have precluded the application of computational approaches in understanding PIEZO1. Attempts to study PIEZO1 are further complicated by the functional redundancy of PIEZO1 in various biological systems. Despite these challenges, the unique LoF and GoF variants position PIEZO1 to be a candidate for evaluation with pathogenicity prediction algorithms which may help decipher variants between these two mutation types. Currently, limited literature exists on the abilities of pathogenicity prediction algorithms to decipher variants of LoF from GoF. Herein, we sought to perform a case study of PIEZO1 variants using these computational approaches.

We performed comparative benchmarking of multiple pathogenicity tools in assessing these mutation types, especially tools utilizing machine-learning based algorithms such as AlphaMissense and ESM-1b (Cheng et al., 2023; Brandes et al., 2023). Previous work has assessed more than 17,000 variants across five genes (*DDX3X, MSH2, PTEN, KCNQ4*, and *BRCA1*) and found that AM scores correlate with *in vitro* functional assay data (Ljungdahl et al., 2024). This correlation has also been observed for variants in the *CFTR* gene implicated in cystic fibrosis and multiple amyloidogenic genes (*APP*, *PSEN1*, and *PSEN2*) (McDonald et al., 2024; Pillai et al., 2025). Our previous work has established pathogenicity prediction tools that are capable of assessing variants occurring on intrinsically disordered regions of proteins, allowing for analysis of variants located on unresolved residues of the PIEZO1 structure (Pillai et al., 2025). Missense mutations demonstrate broad diversity in functional biological impacts, including degrees of LoF, risk-imposing, benign or protective, and GoF variants. Interestingly, Ljungdahl et al., (2023) found that both LoF and GoF variants received high pathogenicity scoring for the *KCNQ4* gene. A potential hypothesis for these approaches identifying GoF with high pathogenicity is that they impart significant changes to the protein structure, resulting in the introduction of novel biological functions, and thereby being predicted as pathogenic. For instance, GoF mutations (e.g. G12V, G12C, G12D) in the *KRAS* oncogene result in either strong stabilization or destabilization of free energy of the protein structure as shown in molecular dynamic simulations and deep mutational scanning experiments (Pandey et al., 2025; Kwon et al., 2024). Yet, these variants are incompletely understood. There is a possibility that GoF imposes minimal structural effects but rather affects unstructured regions or other dependent biological factors implicated in protein function (Li et al., 2019; Li et al., 2018). Accurately deciphering LoF from GoF variants may support numerous applications in precision medicine, and the potential for application of predictive *in silico* approaches to answering this question remains uncertain.

Herein, we benchmarked multiple pathogenicity prediction algorithms and calculated the free energy of stability data for 75 unique missense variants implicated in seven reported clinical phenotypes linked to PIEZO1, including LoF mutations in lymphatic dysplasia and GoF in HX. This is the first case study to our knowledge to directly evaluate the capabilities of pathogenicity prediction algorithms against these mutation types.

## 2. Materials and Methods

### 2.1. 3D Structure, Variants, and Phenotypes of PIEZO1

A comprehensive table of missense variants for PIEZO1 and their reported phenotypes was obtained from More et al., (2020). Of the 75 unique variants, only one, F2458L, has been reported to cause both LD and HX. This variant was included twice as two data points in our analyses. We modified this table to sort by phenotype and obtained the functional and stability data for this list (**Supplementary Material)**. Next, we utilized the predicted 3D structure of PIEZO1 (AF-Q92508-F1-v4) derived from AlphaFold2 (AF2) in the AlphaFold Structure Database (Jumper et al., 2021; Varadi et al., 2022). We reviewed seven *Mus musculus* derived cryo-EM structures from Protein Data Bank (PDB: 6B3R; 7WLT; 7WLU; 8IMZ; 6LQI; 5Z10; 6BPZ; 3JAC). Visualization was completed in PyMOL (Schrödinger LLC, Portland, OR, USA, Version 3.1.0). Additionally, one recently resolved human PIEZO1 structure was reviewed as well (PDB: 8YEZ).

### 2.2. Pathogenicity and Stability Scoring for PIEZO1 Variants

AlphaMissense scores were obtained from the AlphaFold Structure Database (Heatmap), and established classification cutoffs were used. Variants with scores above 0.56 were classified pathogenic, scores below 0.34 were classified benign, and scores between 0.34-0.56 were classified as ambiguous (Cheng et al., 2023). Likewise, CADD scores were obtained from the ProtVar server from the EMBL-EBI. Scores below 15 were classified as likely benign and those above 15 were classified as likely pathogenic (https://www.ebi.ac.uk/ProtVar/) (Rentzsch et al., 2018; Stephenson et al., (2024)). For Evolutionary Scale Modeling-1b (ESM-1b), the negative log-likelihood ratio (-LLR) was used for scoring. A -LLR of less than seven was classified as corresponding to a benign variant, a score greater than eight was classified as corresponding to a pathogenic variant, and a score between seven and eight inclusive was considered ambiguous (https://huggingface.co/spaces/ntranoslab/esm_variants) (Brandes et al., 2023). Lastly, scoring of the Evolutionary Model of Variant Effect (EVE) (https://evemodel.org/) followed the criteria established by Frazer et al. (2021), where scores greater than 0.65 were considered pathogenic, those below 0.35 were considered benign, and those in between 0.35 and 0.65 were considered ambiguous. Next, with the AF2 structure of PIEZO1, we used a modified version of the open-source, peer reviewed protocol defined by Pillai et al., (2025) to calculate the change in free energy (ΔΔG) upon the amino acid substitution in each variant. To evaluate the stability change in the missense variants, we considered using the PremPS, DDMUT, DynaMut2, MAESTRO, DDGEmb, and INPS-MD methods for computation (Chen et al., 2020; Zhou et al., 2023; Rodrigues et al., 2020; Laimer et al., 2015; Savojardo et al., 2016; Savojardo et al., 2024). However, given the size of the ion channel exceeded the amino acid limit (AA < 2500) for most of these computational approaches, we elected to use PremPS which was trained on a balanced dataset of stabilizing and destabilizing missense variants. Lastly, a preliminary ensemble model was produced with Python version 3.10, using libraries including pandas, matplotlib, sklearn, and seaborn (McKinney 2011; Hunter, 2007; Waskom 2021; Pedregosa et al., 2011). All programming occurred on a Google Colab environment and used a T4 GPU.

### 2.3. Statistical analyses

All statistical analyses and data plotting was performed using GraphPad Prism (version 10.3.1) (GraphPad Software, La Jolla, CA, USA). Unless stated otherwise, all data displayed has been reported as the mean ± standard error of the mean (SEM). *p*-values less than 0.05 were deemed statistically significant. When comparing individual variables, a one-sample Student’s *t*-test was used for significance testing. Likewise, when comparing multiple variables, a Mann-Whitney U test was used for evaluating statistical significance. For linear correlational analyses, we calculated Pearson’s correlation coefficients and Spearman’s correlation coefficients to determine whether the predicted slope was significantly non-zero. A GraphPad file containing all statistical analyses and visualizations has been provided in the **Supplementary Material**.

## 3. Results

### 3.1. AlphaFold2 predictions are consistent with cryo-EM structures of wild-type PIEZO1

Before evaluating our selected pathogenicity prediction algorithms, we sought to confirm the accuracy of the AF2 structure (AF-Q92508-F1-v4) of PIEZO1 as a prerequisite to measuring the change in free energies in unresolved regions of the protein in point mutation variants. At the time of writing, there were no structures in the Protein Data Bank with a complete resolution of all PIEZO1 domains. Therefore, we evaluated the structure of the AF2 prediction against all partially resolved cryo-EM structures of PIEZO1, described in Section 3.2. When superimposing the AF2 structure with each individual subunit of the homotrimer (*n* = 24), the mean root mean square deviation of atomic position (r.m.s.d.) was found to be 4.350 ± 0.212 Å, demonstrating similarity of the AF2 prediction with partially resolved structures. We utilized the cryo-EM structure (PDB: 5Z10) as a template to assemble a homotrimeric model of PIEZO1 using the AF2 prediction (**Fig 1A**). We also found that the AF2 structure was nearly identical to a human PIEZO1 structure recently resolved by Shan et al., (2024), where the r.m.s.d. was 2.445 ± 0.007 Å (*n* = 3). The overall predicted local distance difference test (pLDDT) score for the structure was 73.44, and those for the 75 variants was 79.91 ± 1.68, both indicating high confidence predictions of the 3D structure. However, there were 13 variants (17.57%) that were located on low confidence residues of the structure, displayed in **Fig 1B**. Given the unique structure of PIEZO1 from other ion channels, the MMseqs2 algorithm used to create multiple-sequence alignment in the AF2 prediction identified evolutionary similarity with varying species, including *Rattus norvegicus*, *Mus musculus*, *Glycine max*, *Arabidopsis thaliana*, and *Zea mays*, and likely will not hinder conservation data utilized for AM predictions. Therefore, with consideration of the structure and sequence similarity along with the pLDDT scores, we began our computational analyses with the AF2 structure.

**Figure 1.**
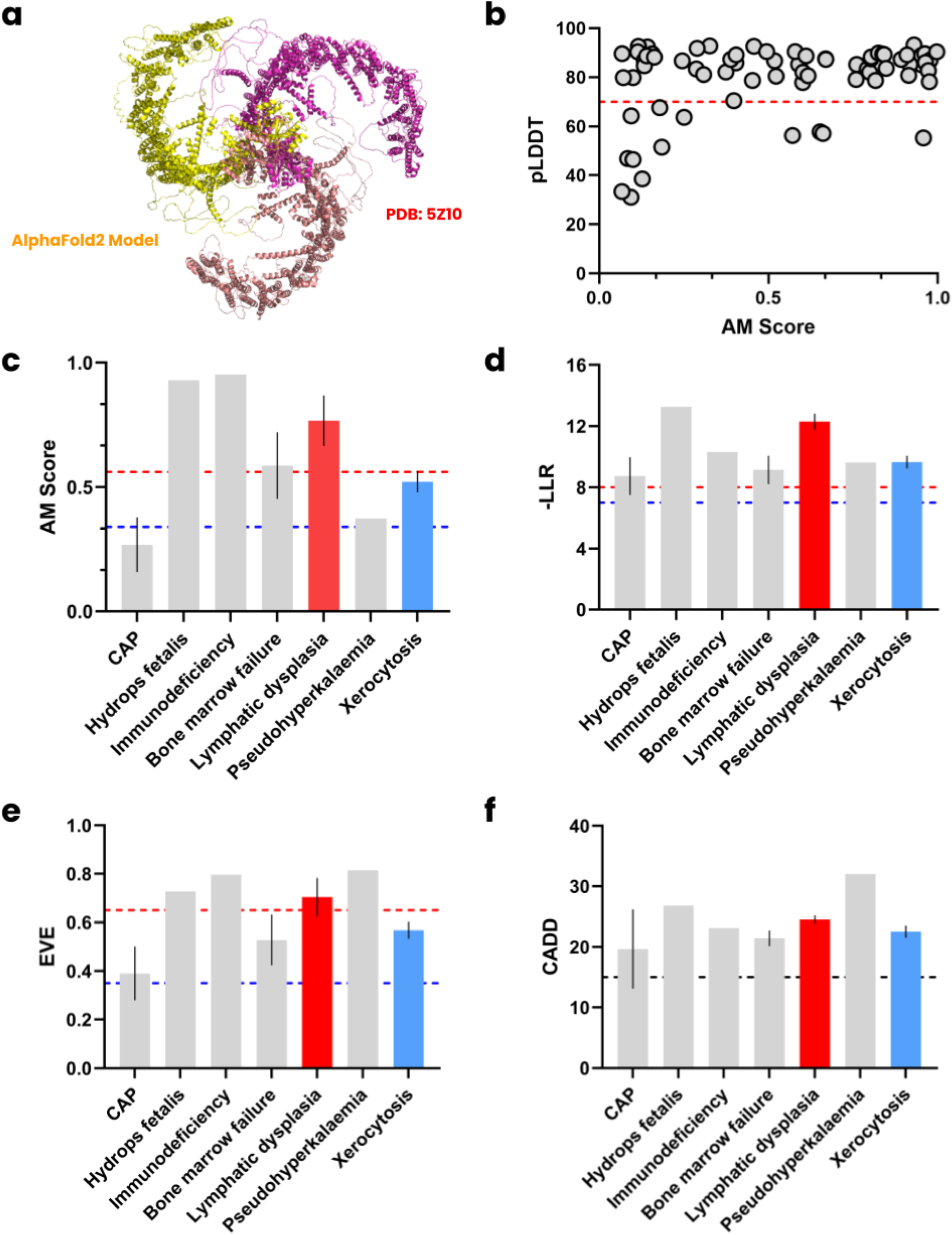
Overall pathogenicity for the phenotypes of PIEZO1 variants. **(a)** AlphaFold2 model of PIEZO1 homotrimer superimposed from a prior cryo-EM structure (PDB: 5Z). **(b)** The pLDDT scores for the 75 missense variants from More et al., (2020). Dotted red line indicates cutoff for high confidence prediction. **(c)** AM scoring for the 7 unique phenotypes of missense variants, where the dotted red and blue line indicates the cutoff for pathogenic and benign classification, respectively. **(d)** The negative log-likelihood ratio from ESM-1b according to phenotype, where the red and blue line indicates the cutoff for pathogenic and benign classification, respectively. **(e)** Distribution of EVE scores, with dotted red and blue lines indicating the cutoff for pathogenic and benign classification, respectively. **(f)** CADD scoring is distributed for the phenotypes with a blue line indicating the benign classification cutoff.

### 3.2. AlphaMissense identifies LoF, but fails with GoF variants in comparison to ESM-1b

Next, we evaluated pathogenicity scoring from AM, ESM-1b, CADD, and EVE for each of the unique phenotypes occurring from the PIEZO1 variants, including bone marrow failure, colorectal adenomatous polyposis (CAP), hydrops fetalis, immunodeficiency, LD, HX, and pseudohyperkalemia. LD is caused by LoF and HX from GoF, respectively. AM scores for all available phenotypes are displayed in **Fig 1C**. The mean score of LD variants was 0.7672 ± 0.1017 (*n* = 6) and that of HX variants was 0.5213 ± 0.0425 (*n* = 56). AM was able to decipher LoF variants with high scores despite a limited sample size for the LD phenotypes, which validates prior studies evaluating AM (McDonald et al., 2024; Ljungdahl et al., 2023; Pillai et al., 2025). Interestingly, the GoF variants for HX were found to be within the range of uncertain classification (AM: 0.34-0.56). Seven variants were classified as uncertain (L2277M, L2192I, K2502R, A2003T, R1797C, A2020T, and E2461K). There were 29 variants classified as pathogenic and 20 as benign. These findings suggest that there is no clear trend of pathogenicity for GoF variants in PIEZO1, contradicting the high pathogenicity consistently predicted by AM with the GoF variants in the *KCNQ4*, a gene that encodes a potassium channel protein (Ljungdahl et al., 2023). While there remains no clear biological trend reported for bone marrow failure and CAP implicated from PIEZO1 variants, both were within range of uncertain pathogenicity at 0.5863 ± 0.1337 (*n* = 6) and 0.2686 ± 0.1101 (*n* = 6), respectively. No clear pathogenicity is observed for hydrops fetalis, immunodeficiency, and pseudohyperkalemia as there remains only a single variant reported in literature. The AM scores at the pathogenic benchmark (0.56) were not significant for both LD (*p* = 0.0972) and HX (*p* = 0.3669). However, AM scores for both LD (*p* = 0.0085) and HX (*p* < 0.0001) were significantly above the benign benchmark (0.34). The non-significance observed for pathogenic LD-causing PIEZO1 variant AM scores may be attributed to the low sample size of variants observed in patients.

The trends of predicted pathogenicity among the phenotypes from ESM-1b is shown in **Fig 1D**. Generally, all phenotypes were determined to be pathogenic (-LLR > 8). The mean -LLR score for LD-causing variants was 12.29 ± 0.53 and that for HX was 9.65 ± 0.41 (*p* = 0.0268). Both LoF (*p* = 0.0005) and GoF variants (*p* = 0.0002) yielded scores significantly greater than the pathogenic threshold (-LLR = 8). The remaining phenotypes were not significantly different from this threshold (*p* > 0.05). These trends appear to align with the expected effects of LoF and GoF variants described in Section 1. The ESM-1b model is unique from AM in that the LLR takes consideration of GoF variants (Brandes et al., 2023), where the negative likelihood is decreased but can still be pathogenic in classification. CAP (*p* = 0.5884) and bone marrow failure (*p* = 0.2733) were also considered uncertain by ESM-1b, while the hydrops fetalis, immunodeficiency, pseudohyperkalemia were also classified above the pathogenic threshold. Overall, deviations among the LoF and GoF variants as predicted by ESM-1b in relation to the clinical ground truth were reduced compared to AM and a difference in likelihood was observed. It appears that while both mutation types are pathogenic, the LoF imparts a greater likelihood between the wild-type and mutant.

Similar trends were also observed with EVE, where LoF variants do not have significantly greater predicted pathogenicity values compared to GoF variants (*p* = 0.2990). Scores for CAP, bone marrow failure, LD, and HX phenotypes were within the range of ambiguity, (**Fig 1E**). CADD exhibited similar results; LoF variant scores were, on average, greater than scores for GoF cairants, though not to a statistically significant degree (*p* = 0.6714). Similarly to our previous reporting with variants on amyloidogenic proteins (Pillai et al., 2025), CADD scores appear to be biased towards pathogenic predictions (**Fig 1F**). With the exception of ESM-1b, LoF variants tend to yield a pathogenic classification to a non-significant but greater degree than by GoF variants across the evaluated pathogenicity prediction algorithms.

### 3.3. GoF PIEZO1 variants are correlated with predicted pathogenicity of EVE and CADD

Correlational analyses were then performed between pathogenicity scores obtained from the four computational approaches and computed changes in free energy upon point mutation using the balanced machine-learning model PremPS (**Fig 2A**). There appeared a statistically insignificant weak negative correlation between the AM predicted pathogenicity scores and resulting stability (*p* = 0.2457). Additionally, there appeared no clear trend between LoF variant scores and PremPS derived free energies as well as between scores for variants of unknown significance (VUS) and PremPS derived free energies. This trend was also observed in ESM-1b calculated variant scores (**Fig 2B**). However, the weak negative correlation was statistically significant for the distribution of EVE (*p* = 0.0276) (**Fig 2C**). Likewise, though the predicted slope was not significantly non-zero for CADD (**Fig 2D**), Spearman’s ⍴ was significantly negatively correlated (*p* = 0.0073). From the individual ΔΔG, HX GoF variants caused significant destabilization to the protein structure at −0.5471 ± 0.0932 kcal/mol (*p* < 0.0001), while LD LoF variants were not significant at 0.0717 ± 0.4931 kcal/mol (*p* = 0.8901), respectively. Remaining VUS variants were found to not cause significant destabilization (**Supplementary Material**).

**Figure 2.**
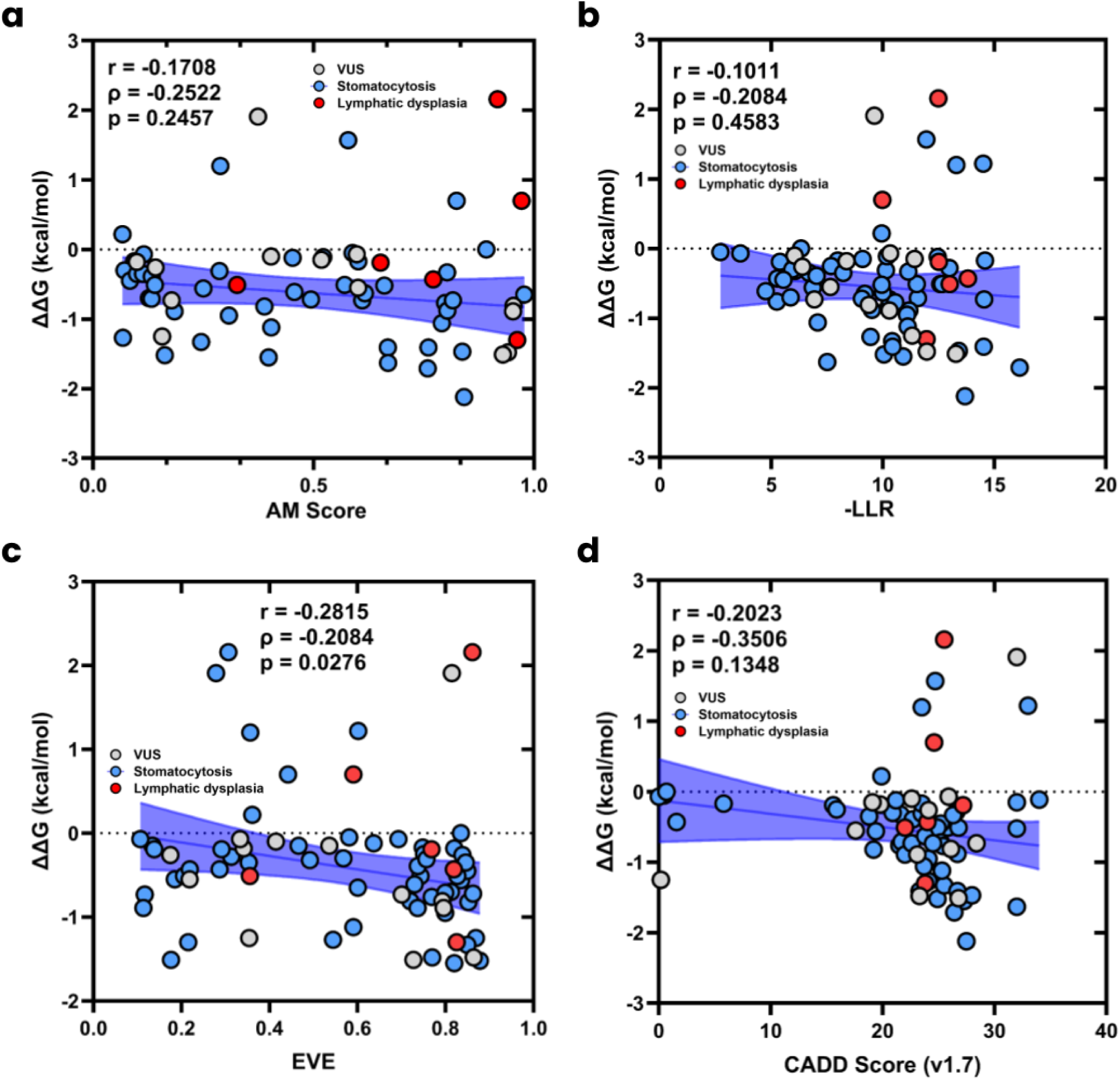
Computed free change upon amino acid substitution. **(a)** Correlation of AM predicted pathogenicity against change in free energy upon mutation, including the legend for each phenotype. **(b)** Correlation of ESM-1b predicted pathogenicity against change in free energy upon mutation, including the legend for each phenotype. **(c)** Correlation of EVE predicted pathogenicity against change in free energy upon mutation, including the legend for each phenotype. **(d)** Correlation of CADD predicted pathogenicity against change in free energy upon mutation, including the legend for each phenotype.

### 3.4. Potential ensemble approach to pathogenicity prediction of PIEZO1 variants

As described in our previous reporting with amyloidogenic genes (Pillai et al., 2025), there appears to be great promise in the possibility of ensemble methods towards accurately predicting the functional effects of missense variants. Indeed, in internal validation studies by Cheng et al., (2023), the REVEL ensemble algorithm performed the second highest in accuracy and precision on the ClinVar database (shown in Fig. 2 of publication), suggesting potential interest in ensemble techniques for deep learning-based approaches. Herein, we developed a preliminary weighted ensemble approach with validation using PIEZO1 variants that have received classification as either pathogenic or benign from the ClinVar dataset (*n* = 118). The in-house python script to this model has been provided in the **Supplementary Material**. Using the scoring from each model without consideration of the thresholds for pathogenic or benign classification, we normalized the data and optimized the weights of individual models dynamically to achieve the highest area under the curve (AUC) using the sequential least square optimization method. We arbitrarily labeled scores from 0.4 to 0.6 as ambiguous, which were deemed incorrect predictions for ROC-AUC analyses. Surprisingly, the weighting that achieved the highest AUC was a balanced incorporation of (0.25) all four algorithm scores per variant. We found that the overall accuracy of the ensemble model was 81.4% with an AUC of 0.955 and generally outperformed all models except for AM which reported the highest accuracy of 86.4% and an AUC of 0.959. Metrics for this preliminary ensemble model have been displayed in **Fig 3A-C**. Overall, these results indicate that AM still retains the best performance of the evaluated approaches and the ensemble model.

**Figure 3.**
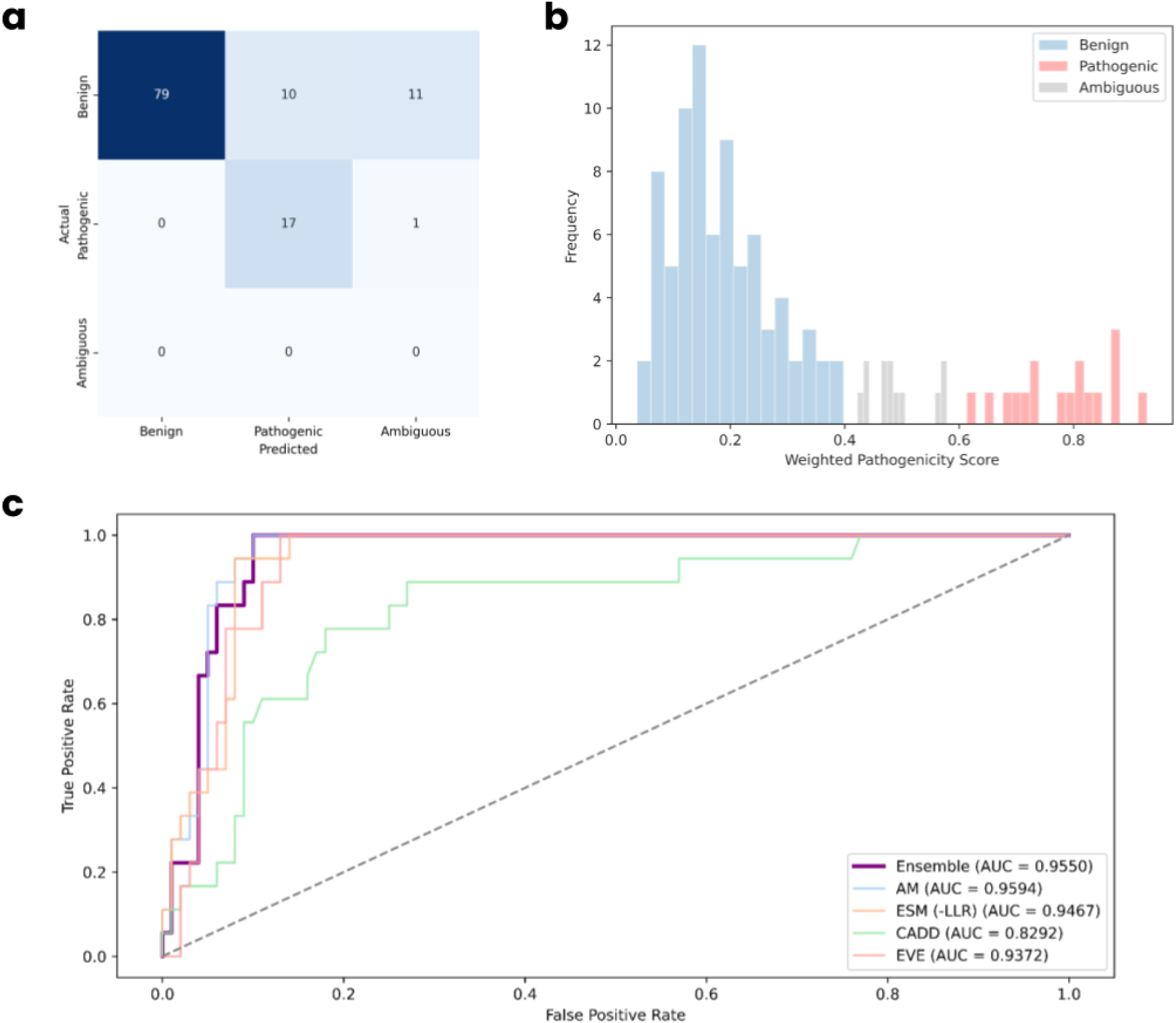
Performance of preliminary ensemble model for PIEZO1 variants. **(a)** Confusion matrix of performance of the model, indicating high accuracy with significant false positives for benign variants. **(b)** Distribution of predicted scores. **(c)** ROC-AUC analyses results of individual models against the ensemble approach.

## 4. Discussion

Classifying the pathogenicity of missense variants is a crucial application in precision medicine, and remains a recurrent challenge due to the large diversity of possible biological effects. Over the past decade, the majority of computational approaches have focused on determining degrees of LoF, but limited literature exists on those for GoF variants. Due to the breakthrough of AF2, many of these approaches have adopted structural data in consideration of these effects, particularly AM. Numerous studies have demonstrated that AM is correlated with *in vitro* functional assays of variants, but no studies have evaluated genes involving both GoF and LoF variants. Therefore, we sought to perform a case study of PIEZO1 variants that involved both mutation types. In this study, we examined the performance of several pathogenicity prediction models in the context of the *PIEZO1* gene and LoF or GoF variants deriving from missense mutations. We found that AM, a deep learning classifier that convolves AlphaFold structural context with evolutionary conservation, is adept at recognizing deleterious LoF variants but is less able to recognize GoF variants (Cheng et al., 2023). AM assigned higher pathogenicity scores to known LoF mutations from PIEZO1 mutation-associated LD, accurately indicating these variants as functionally damaging and reflecting a training emphasis on identifying variants that disrupt protein structure or conservation in LoF mutations. By contrast, AM was more likely to misclassify known GoF mutations causing HX as benign or of uncertain significance.

These results are consistent with the broader literature on PIEZO1 and variant pathogenicity. How mechanical force activates PIEZO1 remains only partially understood (Wang et al., 2018). Diseases caused by *PIEZO1* mutations illustrate two opposite mechanisms. GoF variants typically do not grossly destabilize the protein; instead, they alter channel gating kinetics or regulatory domains. For example, HX-associated PIEZO1 mutations have been shown to slow channel inactivation kinetics (Lukacs et al., 2015), resulting in prolonged cation influx without abolishment of function. Such changes are pathogenic *in vivo* but may not trigger recognition of conventional damage signatures that algorithms like AM recognize. In contrast, LoF mutations often disrupt conserved residues or protein folding, changes that are readily detected by conservation- and structure-based models. This mechanistic discrepancy in variant effects may explain the underperformance of AM on GoF predictions.

Interestingly, the protein language model ESM-1b demonstrated a greater balance in sensitivity to both LoF and GoF variants than AM. Unlike AM, ESM-1b is an unsupervised model trained on approximately 250 million general protein sequence patterns. It has previously been shown that such language models are capable of predicting mutational impacts in a zero-shot manner by identifying the degree to which a mutation appears deleterious or unusual against learned sequence regularities (Meier et al., 2021). ESM-1b’s predictions herein flagged not only the LoF variants but also several GoF variants as outliers, despite only with moderate confidence. This suggests that certain gain-of-function mutations may induce sequence features, for instance at critical motifs such as gating motifs, that the language model recognizes as evolutionarily atypical despite not benign overtly destabilizing. However, ESM-1b was less sensitive than AM for some LoF variants, reflecting a possible trade-off. While previous work has focused primarily on distinguishing pathogenic from benign variants, herein, we directly assessed pathogenicity prediction model performance in identifying distinct pathogenic mechanisms. PIEZO1 was selected for evaluation as LoF and GoF mutations produce variants associated with well-characterized pathophysiology. Our findings underscore a potential limitation in current prediction algorithms in which pathogenicity is predominantly associated with structural disruption-based LoF.

Our study contains several limitations. The limited number of confirmed PIEZO1 LoF mutations restricts statistical robustness and generalizability. Furthermore, despite advancements in cryo-EM, unresolved structural regions may perturb AlphaFold-derived model predictions. Static structure-based methods may also fail to capture minor conformational changes induced by GoF mutations. Furthermore, pathogenicity prediction tools are constrained by a lack of mechanism specification or by heuristic score determination, altogether reducing model interpretability. Future approaches to improving GoF identification in pathogenicity prediction models may include training models on datasets enriched with activating mutations or utilizing multi-task learning frameworks for simultaneously classifying pathogenicity and functional mechanisms. Improved pathogenicity prediction will advance our understanding of variant pathogenicity and facilitate improved diagnostics and therapeutics development.

## Data Availability

The source data has been deposited in the Supplementary Material and GitHub repository: https://github.com/Joshua-Pillai/PIEZO1.

## Acknowledgements

We would like to thank Dr. Shang Ma, Ph.D. from the Children’s Medical Center Research Institute at the University of Texas Southwestern Medical Center at Dallas for technical assistance with reviewing the presented study.

---

**Fig S1.**
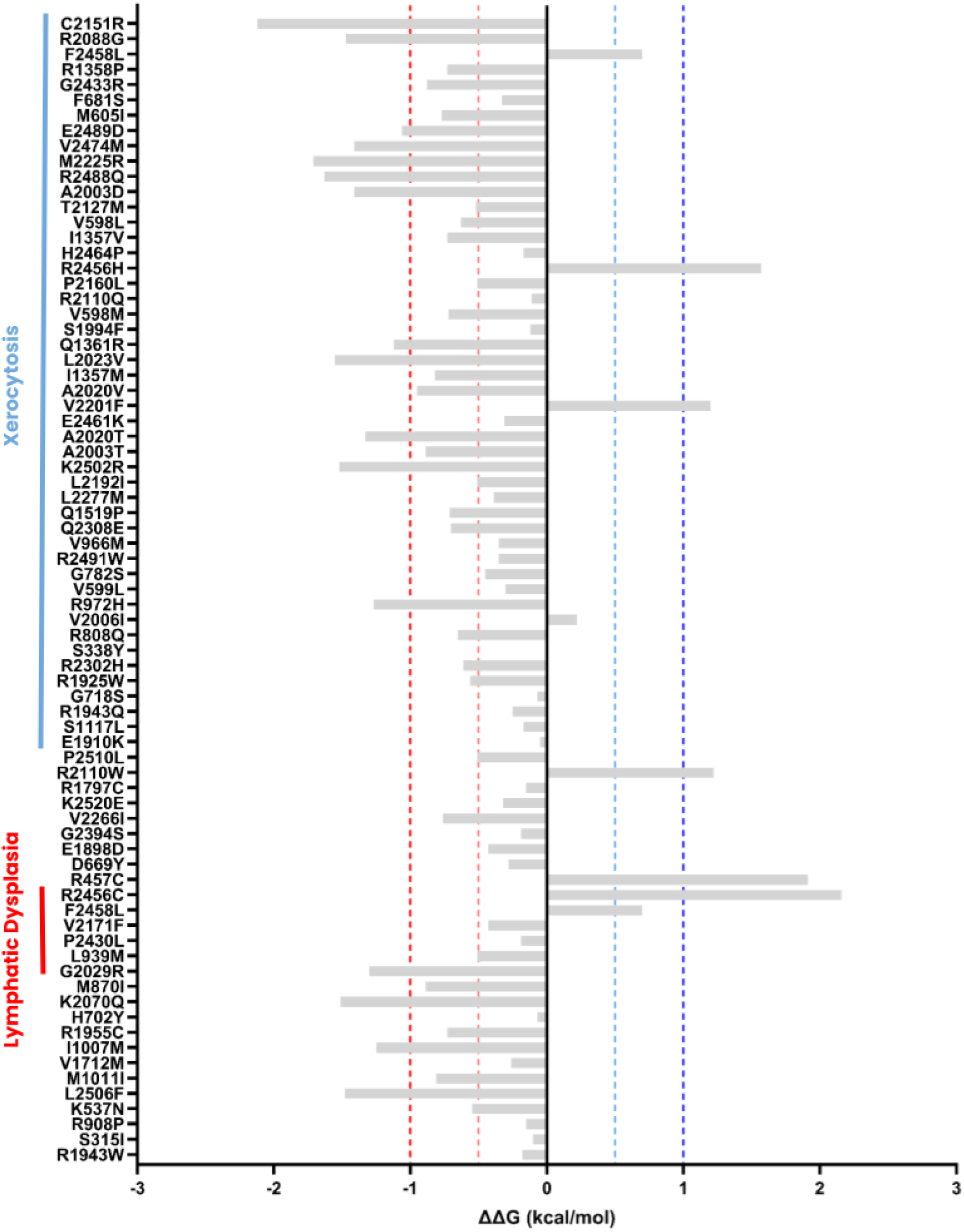

**Supplementary Table 1.** Table of missense variants from More et al., (2020).

**Supplementary Table 2.** The source data of AlphaMissense, CADD, EVE, and ESM-1b prediction scores, along with pLDDT scores.

**Supplementary Table 3.** The source data of change in free energy of stability following the protocol reported by Pillai et al., (2025).

**Supplementary Table 4.** superimposition of cryo-EM and AF2 structures of PIEZO1.

**Supplementary Table 5.** ClinVar dataset for PIEZO1 variants.

**Supplementary Data.** The source files used for statistical analyses and visualization of the reported data in GraphPad Prism.

